# A multi-scale feature fusion model based on biological knowledge graph and transformer-encoder for drug-drug interaction prediction

**DOI:** 10.1101/2024.01.12.575305

**Authors:** Tao Wang, Qiang Deng, Jialu Hu, Yongtian Wang, Jiajie Peng, Jing Chen, Xuequn Shang

## Abstract

Drug-Drug Interaction (DDI) refers to the combined effects that occur when a patient takes multiple medications simultaneously or within the same period. This interaction can either enhance the therapeutic effects of the drugs or inhibit their efficacy, and in severe cases, it can even lead to adverse drug reactions (ADRs). Thus, it is crucial to identify potential DDIs, as this information is significant for both biological research and clinical medicine. However, most existing works only consider the information of individual drugs or focus on the local correlation between a few medical entities, thus overlooking the global performance of the entire human medical system and the potential synergistic effects of multi-scale information. Consequently, these limitations hinder the predictive ability of models. In this paper, we propose an innovative multi-scale feature fusion model called ALG-DDI, which can comprehensively incorporate attribute information, local biological information, and global semantic information. To achieve this, we first employ the Attribute Masking method to obtain the embedding vector of the molecular graph. Next, ALG-DDI leverages heterogeneous graphs to capture the local biological information between drugs and several highly related biological entities. The global semantic information is also learned from the medicine-oriented large knowledge graphs. Finally, we employ a transformer encoder to fuse the multi-scale drug representations and feed the resulting drug pair vector into a fully connected neural network for prediction. Experimental evaluations on datasets of varying sizes and different classification tasks demonstrate that ALG-DDI outperforms other state-of-the-art models.

## Introduction

Drug-Drug Interaction (DDI) refers to the complex effects that occur when a patient takes multiple medications simultaneously or within the same period. These interactions can either enhance the therapeutic effects of drugs or inhibit their efficacy, and in severe cases, they may lead to Adverse Drug Reactions (ADRs). Unintended DDIs are responsible for approximately 3-26% of drug-related hospital admissions [1]. In a 2020 follow-up survey conducted by Laville et al. [2], patients with chronic kidney disease, predominantly aged 69 years old, were found to commonly use five to ten drugs. During the two-year survey, they found that 14.4% of patients had adverse drug reactions, and 32% of them could actually be prevented. These findings underscore the significant implications of drug interactions in practical medical practice and the growing urgency to identify potential DDIs. However, diagnosing DDIs through traditional wet lab experiments for a large number of drug pairs, both in vitro and in vivo, is a time-consuming and costly endeavor. This led to the rising popularity of using computer-based methods, especially machine learning, for DDI prediction. These methods can be roughly classified into three categories: (1) similarity-based methods; (2) graph-based methods; (3) matrix factorization-based methods.

In similarity-based methods, researchers often rely on the drug similarity assumption, where it is believed that if drugs A and B interact to produce a specific biological effect, then drugs similar to drug A (or drug B) are likely to interact with drug B (or drug A) to produce the same effect [3]. Ryu [4] derived a Structural Similarity Profile (SSP) for an individual drug by comparing its ECFPS fingerprint with those of all other drugs, connecting the SSPs of two drugs was then employed as input for a deep neural network to make predictions. Zhang et al. [5] build an ensemble of prediction methods based on 14 kinds of drug–drug similarities from chemical, biological, phenotypic and topological data. Qian et al. [6] utilized feature similarity and feature selection methods to construct a gradient boosting classifier, speeding up the prediction process and achieving robust predictive performance. Gottlieb et al. [7] computed seven types of similarity and combined the two best similarities for each drug pair to generate drug features. Cheng et al. [8] extracted features from Simplified Molecular-Input Line Entry System (SMILES) data and the side effect similarity of drug pairs, applying Support Vector Machines (SVM) to predict DDIs. Rohani et al. [9] calculated the similarity of multiple drugs and Gaussian interaction curves for drug pairs and then selected the most informative and less redundant similarities as features, using the feature vectors of drug pairs as input for neural network predictions.

In recent years, researchers have increasingly focused on graph data. Compared to non-graph-based methods, researchers can leverage the topological information within the graph to enhance node representations, leading to improved performance. Generally, three types of graphs can serve as input: molecular graphs of drugs, representing networks formed by atoms and their connections; DDI networks, derived from known DDI information to infer new DDIs; and networks between drugs and other medical entities. Cami et al. [10] used standard multivariate methods to combine multiple predictor variables for DDI prediction. They constructed a DDI network and extracted several covariates from the network to build logistic regression and linear mixed models. Zitnik et al. [11] developed a graph convolution network encompassing drugs, targets, and side effects, treating DDI prediction as a multi-relational link prediction task. Karim et al. [12] integrated multi-source datasets into a knowledge graph, utilizing knowledge graph embedding methods and convolutional LSTM networks for DDI prediction. Feng et al. [13] employed Graph Convolution Networks (GCN) to extract drug features from the DDI network. Subsequently, they used a Deep Neural Network (DNN) as a predictor to obtain probabilities of interactions between drugs. Wang et al. [14] divided the DDI network into two parts: promoting DDI network and inhibiting DDI network, where the edges denoted the promoting or inhibiting effects between drugs. Subsequently, they independently extracted drug features from these two networks and combined them for prediction.

The DDI prediction task can be represented as a matrix completion task, which aims to predict unobserved interactions [15]. Typical matrix factorization methods include non-negative matrix factorization, singular value decomposition and so on. Besides, some methods develop novel matrix factorization models based on manifold learning algorithms, artificial neural networks and so on. Yu et al. [16] developed a novel method named DDINMF, based on semi-non-negative matrix factorization. Zhang et al. [5] proposed a manifold-regularized matrix factorization approach for DDI prediction. Zhu et al. [17] devised a dependency network to simulate the relationships between drugs and introduced a probabilistic dependency matrix three-factorization method, known as Property-supervised learning Model Probability Dependency Matrix Three-Factorization (PDMTF), for DDI prediction.

Generally, most previous works utilized large labeled data and merely considered the structure or sequence information of drugs, or only focused on the local correlations between drugs and certain key medical entities(e.g. protein, disease and side effect), while lacking a comprehensive consideration of the global scale of drug interactions in human medical system.This, to some extent, constrained the model’s performance. To address the limitations, we propose a novel multi-scale feature fusion model named ALG-DDI, which is a deep learning framework for DDI prediction and DDI events prediction. Specifically, we first extracted drug attribute information, local information, and global information from the molecular graph of drugs, the heterogeneous graph between drugs and key medical entities, and the medicine-oriented knowledge graph, respectively. Subsequently, to integrate information from different scales, we utilized a Transformer encoder for feature fusion. Through self-attention mechanisms, ALG-DDI can explore synergistic effects between different scales, obtaining high-quality drug embeddings. Moreover,we evaluated the ALG-DDI model on three datasets of different scales, the experimental results demonstrate that ALG-DDI achieved the best performance, thus supporting the effectiveness of the idea of multi-scale feature fusion. The main contributions of this article can be summarized as follows:

- Based on the idea of multi-scale feature fusion, we propose an ALG-DDI model that combines medicine-oriented knowledge graphs and attention mechanisms for DDI prediction and DDI events prediction. It can effectively extract and integrate the features from drug molecular graph, local correlation and knowledge graph.
- We use a feature fusion strategy based on Transformer encoder, which can dynamically assign weights to three different scales of drug information through self attention mechanism, ultimately obtaining a comprehensive representation. Compared to other common fusion methods, our strategy can achieve better performance.
- We compared ALG-DDI with several state-of-art works and variants of our model for ablation study. Experimental results have shown that our work outperformed the baselines on three different datasets.

## Materials and Methodology

### Dataset description

DrugBank is a comprehensive online database that provides extensive biochemical and pharmacological information about drugs, including details about their mechanisms and targets [18]. In this study, We retrieved the most up-to-date version of the structure external links dataset for FDA (Food and Drug Administration) drugs from Drugbank. It includes valuable structure information in the form of InchI/InchI Key/SMILES, as well as identifiers for other drug structure resources, such as ChEBI, ChEMBL, etc. The primary use of this dataset is to obtain SMILES sequences of drugs and filter the DDI data, ensuring that only FDA-approved drugs are included in our experiment. To test the predictive performance and robustness of our model across datasets of different sizes, we utilized three DDI datasets for training and evaluation purposes. The first dataset, named as DS1, was extracted from DrugBank 4.0, which was released in 2014 [18]. After filtering, we obtained 1,294 drugs and 80,866 DDIs from this dataset. The second dataset, named DS2, is derived from DrugBank 5.0, released in 2018, containing nearly six times the amount of DDI data compared to version 4.0 [19]. By applying the same filtering operation, we obtained 1,637 drugs and 392,553 DDIs for DS2. Lastly, the third dataset, DS3, was sourced from PrimeKG, a large-scale medical knowledge graph published by Chandak et al. [20] in 2022. This dataset integrates 20 high-quality resources, encompassing 17,080 issues with 4,050,249 relationships across ten major biological scales. Notably, DS3 includes an extensive number of connecting edges between drugs and other medical entities. To ensure no label leakage during the training process, we removed the edges between drugs, resulting in a total of 1,202,514 triplet data for DS3. For additional information, please refer to Table 1.

**Table 1.**
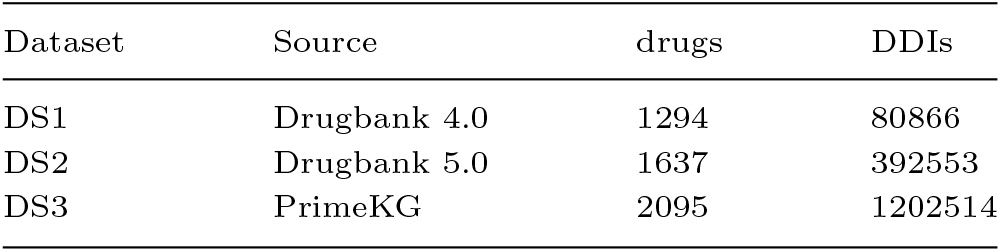
The details of datasets.

| Dataset | Source | drugs | DDIs |
| --- | --- | --- | --- |
| DS1 | Drugbank 4.0 | 1294 | 80866 |
| DS2 | Drugbank 5.0 | 1637 | 392553 |
| DS3 | PrimeKG | 2095 | 1202514 |

### Overview of ALG-DDI

The ALG-DDI framework is presented Figure 1 to illustrate its structure. Our framework consists of three distinct modules: the multi-scale drug feature extraction module I, the feature fusion module II, and the prediction module III. In module I, we adopt a method that combines Attribute Masking and Average Pooling to extract drug attribute information (AI) from molecular graphs, as depicted in I-a. Subsequently, we construct heterogeneous graphs for drug-side effects, drug-disease, and drug-protein interactions, Through heterogeneous graph learning and data alignment, we obtain drug local information (LI), as shown in I-b. Finally,using the ComplEx knowledge graph embedding model, we acquire entity embeddings in PrimeKG, enabling the extraction of drug semantic features as global information (GI), as shown in I-c.In module II, we employ a Transformer encoder that utilizes self-attention mechanisms to fuse drug information from the three mentioned scales. This process allows us to obtain a high-quality and expressive final representation of drugs. In module III, we concatenate the representations of two drugs within a drug pair to create a feature vector. We utilize a three-layer fully connected neural network to make predictions about potential DDIs. The following section provides a detailed explanation of our framework.

**Fig. 1.**
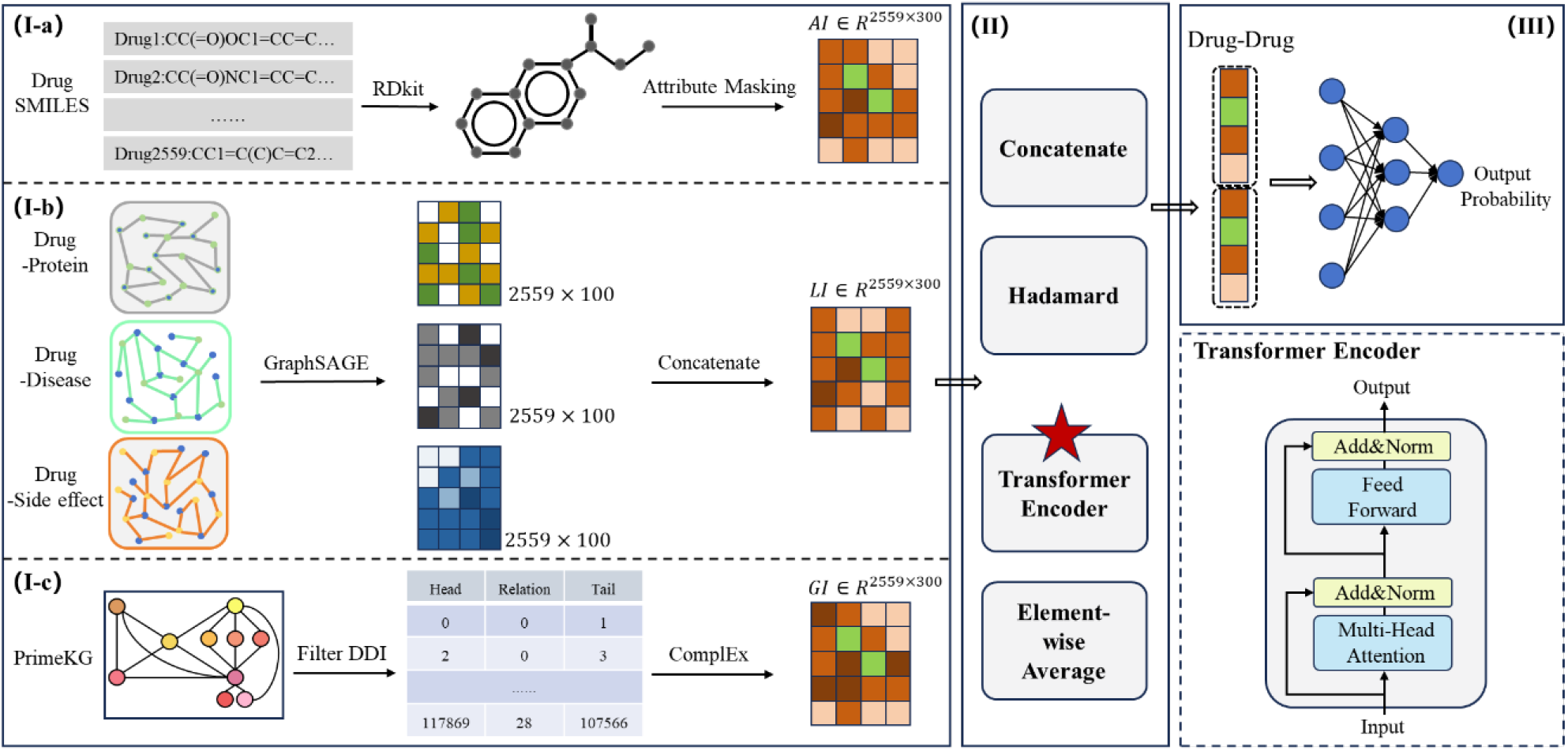
ALG-DDI workflow

### Extraction of multi-scale drug feature

#### Attribute information representation

For each FDA-approved drug *d*_*i*_ ∈ *D*, we convert it into the corresponding molecular graph *g*_*i*_ = (*V, E*) using the RDkit toolkit [21] based on its SMILES string. Here, *V* represents the set of nodes in the molecular graph, which correspond to atoms, and *E* represents the set of edges, which are actually the chemical bonds. To obtain a representation of the molecular graph as the attribute information for the drug, we utilize a pre-trained model called Attribute Masking, proposed by Hu et al [22]. In this model, each node in the molecular graph is embedded using Attribute Masking, and the embeddings of all nodes are then averaged to obtain the representation. Specifically, in the first stage, we use molecular graph, raw node features, and edge features as inputs to calculate the embedding for each atom using Attribute Masking. This method involves setting the features of certain nodes (i.e., masked nodes) to special values and iteratively updating the embeddings of all nodes through a Graph Neural Network (GNN) [23]. The iterative process of node embedding can be represented as follows:

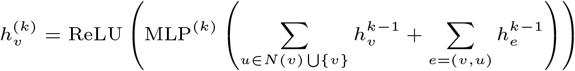

where *k* denotes the index of GNN layers, *N (v)* represents a set of nodes that are adjacent to *v*, and *e* = (*v, v*) signifies the self-loop edge. Additionally, to enhance the representation of the masked node, an embedding-based linear model is employed, which predicts the masked node attribute and aims to preserve the original attributes as closely as possible. In the second stage, we utilize average pooling to aggregate the embeddings of all atoms in the molecular graph. This pooled representation serves as an informative summary of the drug’s attribute information. The second stage can be precisely described as follows:

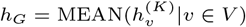

where *K* represents the final layer(i.e. *k* = *K*), and *G* is a set of nodes.Therefore, for FDA drug *d*_*i*_, we can regard *h*_*G*_ as its attribute information *a*_*i*_ *∈* ℝ^*d*^.

#### Local information representation

We consider that the performance of DDI prediction tasks is not only related to the attribute of the drugs themselves but also closely associated with the connections between drugs and other crucial medical entities, which can be used to augment locally detailed information and enhance the capacity of feature learning expression [24]. Therefore, we constructed heterogeneous networks for drug-side effect, drug-protein, drug-disease, and then applied heterogeneous GraphSAGE [25] to capture local information about FDA drugs.

Inspired by RGCN [26], we consider each type of relationship in the heterogeneous graph as a homogeneous graph, the representation learning results of the heterogeneous graph can be translated into the fusion of results from multiple homogeneous graph. For each relationship, we employed GraphSAGE for information propagation. Specifically:

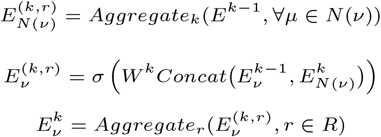

where *k* represents the depth of aggregation, *r* represents the type of relations, *N* (*v*) denotes the set of neighboring nodes for the target node *v, µ* is one of the neighboring nodes, the *Aggregate*_*k*_ operation can be either averaging or multiplication (averaging is used here), *σ* is the activation function (*sigmoid* function is used here), and *W* ^*k*^ represents the weight matrix of the neural network. *Aggregate*_*r*_ operation can be either *sum,mean,min,max* or *mul* in PyG library [27] (*sum* is used here). As *k* increases, GraphSAGE is capable of aggregating information from more layers of neighbors for each node.Finally, for FDA drug *d*_*i*_, we can regard 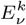 as its local information *l*_*i*_ ∈ ℝ^*d*^.

### Global information representation

The global information of drugs plays a crucial role in our DDI prediction task. The medicine-oriented knowledge graph is pivotal in extracting comprehensive features of drugs, as it encompasses multiple associations between drugs and many biological entities rather than just a few. Compared to the attribute information and the local information, knowledge graphs focus more on the global scale, which can partially overcome the issue of insufficient drug feature expressiveness.

Knowledge graph is a structured way to capture relationships between different entities using sets of ordered triples in the form (head, relationship, tail), which can be defined as follows: G_KG_ = {(*h, r, t*)| *h, t* ∈ *E, r* ∈ *R*}, where *E* represents the set of entities, and *R* represents the set of relationships between entities in *E*. Inspires by Ren et al. [24], we use the effective knowledge graph embedding(KGE) method of ComplEx, proposed by Trouillon et al. [28] to obtain the embedding of each entity. ComplEx obtains the knowledge graph embedding by low-rank decomposition to the knowledge base tensor, which is one of relation adjacent matrixes in knowledge base and the decomposed matrix is used to calculate score. The scoring function *ϕ*(*·*) in ComplEx can be defined as follows:

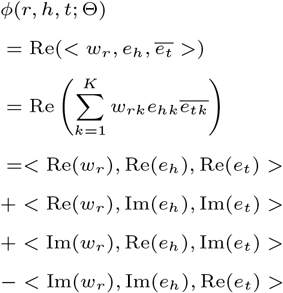

where Θ denotes the parameters of the corresponding model, namely, embedding *w*_*r*_, *e*_*h*_, 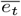, *K* represents a hyperparameter that denotes the embedding dimension, Re(*x*) stands for the part of real vector component of vector *x*, Im(*x*) represents the part of imaginary vector component of vector *x* and 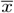 indicates the conjugate vector of *x*, say 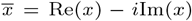. *< · >* means the Hermitian product, which is dot product with complex numbers. If there is a relationship between entity *h*_*i*_ and entity *t*_*i*_, than the log-odd of the probability is:

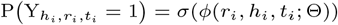

where *σ* represents the sigmoid function and Y_*hi*_,*r*_*i*_,*t*_*i*_ = {−1, 1} indicates the true or false interaction fact of a triple where -1 means no interaction otherwise exists interaction.The ComplEx model is trained to minimize the loss function, described as follows:

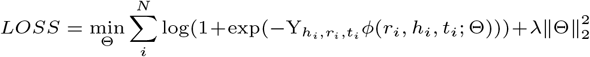

where the *λ* is a hyperparameter. Finally, for FDA drug *d*_*i*_, we can obtain its global information *g*_*i*_ ∈ ℝ^*d*^ through ComplEx model.

### Strategies of feature fusion

In the feature fusion module, we employee a Transformer encoder based on the attention mechanism to fuse three different scales of drug features. The multi-scale drug feature fusion can provide deep learning models with diverse information from different views to accurately predict DDIs, compared to only focusing on one perspective. A Transformer encoder contains multiple Transformer units, while one unit includes self-attention layer, layer normalization, residual connections and feed-forward layer.

### Self-attention mechanism

The self-attention mechanism is the central component within the Transformer encoder. It is a very suitable fusion strategy for our model because it can effectively capture correlations between distinct features, enabling the model to assign suitable weights to each scale, thus enhancing feature fusion.In another word, self-attention mechanism can help us to recognize which scale of features are more important for prediction [15].Multi-head attention, on the other hand, is an extension of self-attention where the mechanism is applied multiple times in parallel, with each application focusing on different aspects of the input. Each “head” captures different patterns and dependencies, providing a richer representation of the input.We denote the representations of drug *d*_*i*_ from three different scales as vectors *a*_*i*_ ∈ ℝ^*d*^, *l*_*i*_ ∈ ℝ^*d*^, *g*_*i*_ ∈ ℝ^*d*^, respectively.

Different scales have inconsistent contributions to the DDI prediction task. Therefore, we concatenate *a*_*i*_, *l*_*i*_, *g*_*i*_ together as input of the Transformer encoder, and use the self-attention mechanism to learn the weight distribution of features at different scales. The multi-head attention is calculated by following formulas.

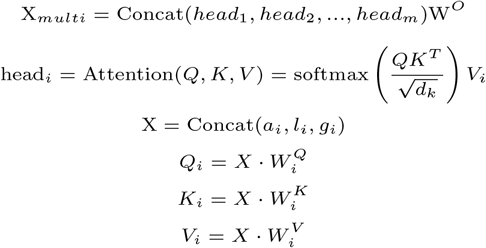

where *X*_*multi*_ ∈ ℝ^*d*^ denotes the fused representation of *d*_*i*_ after transformer encoder, *W* ^*O*^ is a matrix used to transform the concatenated results. For the 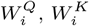 and 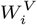 are on behalf of three adaptive weights matrices. *Q*_*i*_, *K*_*i*_ and *V*_*i*_ are the *Q* (Query), *K* (Key) and *V* (Value) matrices derived from the linear transformation of *X*, respectively.

### Residual connections & layer normalization

Residual connections [29], also known as skip connections, are a critical component of the Transformer encoder. They allow the network to skip one or more layers in order to mitigate the vanishing gradient problem, which is common in deep neural networks. In a residual connection, the input to a layer is directly added to the output of the same layer, effectively creating a shortcut path. This helps in the flow of gradients during training and enables the network to learn more efficiently.

Layer normalization [30] is a technique used in deep learning to stabilize training by normalizing the inputs to each layer. In the Transformer encoder, it was used on two occasions: after the self-attention layer and after the feed-forward network layer.

### Comparison with other fusion strategies

In addition to the Transformer encoder, we also employee three common feature fusion methods, including concatenate operation, Hadamard product, and element-wise average, for comparison. These strategies can be described as follows:

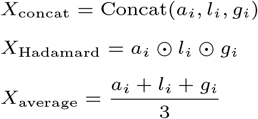

where *X*_*concat*_ ∈ ℝ^3*d*^, *X*_*Hadamard*_ ∈ ℝ*d* and *X*_*average*_ ∈ ℝ^*d*^ denotes the fused representation of *d*_*i*_. It should be noted that due to the inconsistent feature dimensions obtained by the four methods, we use a linear layer to convert fused representation into the final d rug r epresentation *X*_*final*_ with the same dimension *d*.

### Predicting DDIs

In the prediction module, we concatenate the representations of a pair of drugs and then fed them into fully connected layers to predict the DDI probability score. We represent the *X*_*final*_ of drug *d*_*i*_ and drug *d*_*j*_ as *x*_*i*_ and *x*_*j*_, respectively. Then the probability score can be calculated as follows:

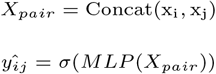

where *y*_*ij*_ is the possibility of drug pair interaction, *σ* is the sigmoid function. During the model training process, we optimize the ALG-DDI parameters by minimizing the cross-entropy loss, as described below:

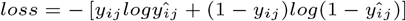

where *y*_*ij*_ denotes the true interaction label for drug pair (*d*_*i*_, *d*_*j*_) in training set.

Strategies for DDI events prediction

To validate the effectiveness of the model in extracting drug features and its generalizability across different scenarios, we further applied ALG-DDI to the DDI events prediction task.The objective of the DDI prediction task is to predict the presence or absence of a relationship between two drugs, while the DDI events prediction task aims to predict the specific relationship. Clearly, the latter holds greater practical value and is more complex than the DDI binary classification task.

Specifically, We obtained the DDI multi-class dataset from the relevant work of Ryu [4], comprising a total of 86 DDI types and 192,284 DDIs. By ensuring that the drugs within the DDI pair are all FDA approved and filtering out categories with insufficient data (fewer than 10 DDIs), we ultimately obtained the multi-class dataset Multi-DS for the DDI events prediction task, which includes 82 categories and a total of 172,323 DDIs. Nevertheless, there is still an imbalance within Multi-DS, with variations in quantity between different categories possibly reaching hundreds or thousands of times. In an effort to mitigate the potential impact of data imbalance on experimental results, we employed the following strategies:

Firstly, we employed a combination of Focal Loss [31] (FL) and Cross Entropy Loss (CE) as our loss function. FL can address the issues of imbalanced sample quantities and difficulty in imbalanced classification. However, due to the instability of FL during training, the neural network requires more training iterations to converge, and in some cases, it may not converge at all [15]. Therefore, in the first half of epochs, we continued to use CE as our loss function, and in the later epochs, we switched to FL. The details are described as follows:

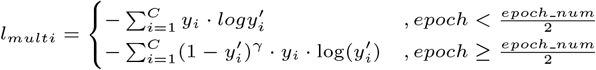

where *C* represents the number of DDI categories, 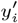 is the predicted probability for the i-th category, *y*_*i*_ is the one-hot value for the i-th category of the true label, and *γ* is the focal parameter in FL, which is used to adjust the weights for easily-classified samples and difficultly-classified samples, aiming to place more emphasis on challenging samples.

In addition, we employed stratified K-fold cross-validation instead of the conventional K-fold cross-validation in DDI events prediction task. This approach ensures that the data proportions within each fold are consistent, avoiding the possibility of sampling all instances of a particular class into the same fold. Stratified K-fold cross-validation is particularly useful when dealing with imbalanced data or datasets with limited volume.

## Experimental Results and Discussion

### Baseline

To evaluate the effectiveness of our model, we compared ALG-DDI with the following state-of-art works.

- **GCN-BMP [32]** utilizes the GCN with Bond-aware Message Propagation to encode molecular graphs. And introduce the self-contained attention mechanism to identify crucial local atoms that align with domain knowledge, providing a level of interpretability.
- **EPGCN-DS [33]**proposed a GCN based framework for type-specific DDI identification from molecular structures. The proposed framework includes an encoder with GCN layers and a decoder that captures complicated interactions while preserves permutation invariant on inputs.
- **MR-GNN [34]** is an end-to-end GNN that employs a multi-resolution-based architecture to extract node features from various neighborhoods of each node. Additionally, it utilizes long short-term memory networks (LSTMs) to summarize local features of each graph and extract interaction features between pairwise graphs
- **DeepDrug [35]**developed a deep learning framework,which can use graph convolutional networks(GCN) to learn graphical representations of drugs and proteins, such as molecular fingerprints and residual structures, to improve prediction accuracy.
- **SSI-DDI [36]**proposed a deep learning framework, which operates directly on the raw molecular graph representations of drugs for richer feature extraction and decomposes the DDI prediction task between two drugs into the identification of pairwise interactions between the substructures of the respective drugs.
- **DeepDDI [4]** designed a multi-class DDI prediction model utilizing a DNN framework. Structural information of each drug in the input drug pair was employed to generate a feature vector referred to as the structural similarity profile, which was designed to effectively capture a drug’s unique structural characteristics.
- **DDIMDL [37]** introduces a multimodal deep learning framework for predicting DDI-associated events. It first constructs deep neural network sub-models based on four types of drug features, and then combines these sub-models to learn cross-modal representations between drug pairs.
- **Lee et al. [38]** utilizes autoencoders and a deep feedforward network, trained on the structural similarity profiles, Gene Ontology term similarity profiles, and target gene similarity profiles of known drug pairs, to predict the pharmacological effects of DDIs.

Among them, GCN-BMP, EPGCN-DS, MR-GNN, SSI-DDI, DeepDrug, and DeepDDI serve as baselines for the DDI prediction task, while DDIMDL, Lee, and DeepDDI serve as baselines for the DDI events prediction task.

### Evaluation metrics

In this work, we employed commonly used classification metrics to evaluate the performance of our framework from various angles, encompassing five key indicators: Accuracy, Precision, Recall, F1, ROC-AUC, PR-AUC. In addition, we also evaluated the performance of ALG-DDI on the DDI events prediction task, so we introduced metrics suitable for multi-class tasks: macro-P (macro-precision), macro-R (macro-recall), and micro-F1 (micro-precision and recall).These evaluation criteria are defined as follows:

- **Accuracy**:accuracy evaluates overall correctness and it can be calculated using the following formula:*Accuracy* = (*TP* +*TN*)*/*(*TP* +*FN* +*FP* +*TN*), where *TP, TN, FP* and *FN* indicate the true positive, true negative, false positive and false negative, respectively.
- **Precision** measures how many of the samples predicted as positive by the model are actually true positive cases, as opposed to being false positives. It can be calculated using the following formula:*Precision* = *TP/*(*TP* + *FP*).
- **Macro-precision** is calculated in multi-class classification tasks by computing the precision for each class and then taking the average of these precision values: 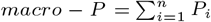
- **Recall**, also known as sensitivity, measures how many of the actual positive cases the model correctly identifies. It can be calculated using the following formula:*Recall* = *TP/*(*TP* + *FN*).
- **Macro-recall** is calculated in multi-class classification tasks by computing the recall for each class and then taking the average of these recall values: 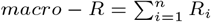
- **F1 score** is a trade-off between precision and recall. And the harmonic mean of average precision and recall score can be calculated using the following formula:*F*_1_ = (2 *\*Precision* Recall*)*/*(*Precision* + *Recall*). Similarly, *macro − F*_1_ can be calculated using the following formula:*macro − F*_1_ = (2 *\* macro − P * macro − R*)*/*(*macro − P* + *macro − R*)
- **ROC-AUC** is the abbreviation for “Area under the Receiver Operating Characteristic Curve”, which represents the area under the plot of the true positive rate against the false positive rate at various thresholds.
- **PR-AUC**: is the abbreviation for “Area under the Precision-Recall Curve”, which represents the area under the plot of the precision rate against recall rate at various thresholds.

### Comparison of other methods

To comprehensively evaluate the performance of ALG-DDI, we compared it with the aforementioned baselines. To ensure fairness, we uniformly utilized the DS2 dataset and employee average values based on 5-fold cross-validation as the experimental results, which are shown in Table 2. The comparison results indicate that ALG-DDI performs best with outstanding improvements of 3.87%-24.45%, 2.74%-27.07%, 4.37%-24.78%, 3.90%-24.37%, 1.38%-18.14%, 1.53%-21.57%, 0.0107-0.2136, 0.0123-0.1365 against others in terms of accuracy, precision, recall, F1, ROC-AUC and PR-AUC, respectively.

**Table 2.** Results of comparing with state-of-the-art method on DS2.

| Method | Accuracy | Precision | Recall | F1 | ROC-AUC | PR-AUC |
| --- | --- | --- | --- | --- | --- | --- |
| GCN-BMP | 0.7422 | 0.7409 | 0.7471 | 0.7431 | 0.8172 | 0.7828 |
| EPGCN-DS | 0.7463 | 0.7081 | 0.8394 | 0.7679 | 0.8224 | 0.7885 |
| MR-GNN | 0.8444 | 0.8191 | 0.8865 | 0.8503 | 0.9185 | 0.8966 |
| SSI-DDI | 0.8612 | 0.8420 | 0.8910 | 0.8651 | 0.9327 | 0.9158 |
| DeepDrug | 0.9022 | 0.8664 | 0.9512 | 0.9068 | 0.9557 | 0.9418 |
| DeepDDI | 0.9480 | 0.9514 | 0.9442 | 0.9478 | 0.9848 | 0.9832 |
| ALG-DDI | **0.9867** | **0.9788** | **0.9949** | **0.9868** | **0.9986** | **0.9985** |

We attribute this superior performance to two factors. First, compared to baseline methods that only focus on a single scale, ALG-DDI has a more powerful capability to exploit multi-scale drug information, including the attribute information of drugs themselves, local information from the interactions with key medical entities, and global information from the large-scale knowledge graph. Second, the Transformer encoder based on the self-attention mechanism effectively integrates multi-scale information by assigning dynamic weights, which contributes to its performance.

### Comparison of different fusion strategies

In the feature fusion module of ALG-DDI, we employee the Transformer encoder as a fusion strategy, achieving notable results. To highlight the effectiveness and superiority of the Transformer encoder in our work, we compared it with four common fusion strategies mentioned above. We conducted 5-fold cross-validation experiments on DS1, DS2, and DS3, and the average experimental results are presented in Table 3.

**Table 3.** The results of different fusion strategies on different datasets.

| Dataset | Method | Accuracy | Precision | Recall | F1 | ROC-AUC | PR-AUC |
| --- | --- | --- | --- | --- | --- | --- | --- |
| DS1 | Concat | 0.9603 | 0.9521 | 0.9695 | 0.9606 | 0.9891 | 0.9852 |
|  | Hadamard | 0.9177 | 0.9034 | 0.9356 | 0.9191 | 0.9670 | 0.9580 |
|  | Average | 0.9474 | 0.9321 | 0.9651 | 0.9483 | 0.9827 | 0.9766 |
|  | Transformer | **0.9681** | **0.9521** | **0.9857** | **0.9686** | **0.9899** | **0.9855** |
| DS2 | Concat | 0.9813 | 0.9736 | 0.9894 | 0.9814 | 0.9977 | 0.9974 |
|  | Hadamard | 0.9197 | 0.9122 | 0.9289 | 0.9204 | 0.9721 | 0.9694 |
|  | Average | 0.9569 | 0.9383 | 0.9781 | 0.9578 | 0.9898 | 0.9879 |
|  | Transformer | **0.9867** | **0.9788** | **0.9949** | **0.9868** | **0.9986** | **0.9985** |
| DS3 | Concat | 0.9756 | 0.9689 | 0.9828 | 0.9758 | 0.9975 | 0.9974 |
|  | Hadamard | 0.8370 | 0.8328 | 0.8434 | 0.8380 | 0.9194 | 0.9166 |
|  | Average | 0.9181 | 0.8956 | 0.9465 | 0.9203 | 0.9774 | 0.9762 |
|  | Transformer | **0.9859** | **0.9800** | **0.9920** | **0.9859** | **0.9991** | **0.9990** |

Firstly, we can observe that ALG-DDI demonstrates good robustness, achieving excellent performance on datasets of various sizes. As the dataset size increases, the results improve even further. Even in DS3, AUROC and AUPR reach the highest values of 0.9991 and 0.9990, respectively. This demonstrates that richer information and more training data enable ALG-DDI to learn more potential patterns, contributing to the improvement of its generalization ability.

Additionally, we observed that as the dataset size increases, the advantage of the Transformer encoder becomes more pronounced. The lead in AUROC grows from 0.0008-0.0229 to 0.0016-0.0797. We attribute this to the self-attention mechanism’s ability to allocate weights to different scale features, enabling it to extract more crucial patterns for the task from large-scale data compared to other methods. Surprisingly, we found that the simplest concatenate operation also yields good results. We attribute this to the fact that drug embeddings have already integrated sufficient rich information through the feature extraction module, making straightforward fusion strategies effective as well.

### Version validation experiment

To further test the model’s effectiveness, we conducted a version validation experiment. We trained the model on a lower version of the dataset and made predictions for all other possible DDIs. We select the top-*N* DDIs based on these scores and verified their existence in higher versions of the dataset. The results are shown in Figure 2. Specifically, we initially compile all possible drug pairs formed by FDA-approved drugs. The DDIs within DS1 are used as a training set, while the remaining DDIs are employed to generate prediction scores. We select DDIs with prediction scores in the top 50, top 100, top 0.1%(7,251 DDIs), and top 0.5%(36,452 DDIs), and examine whether they exist in DS2 and DS3.

**Fig. 2.**
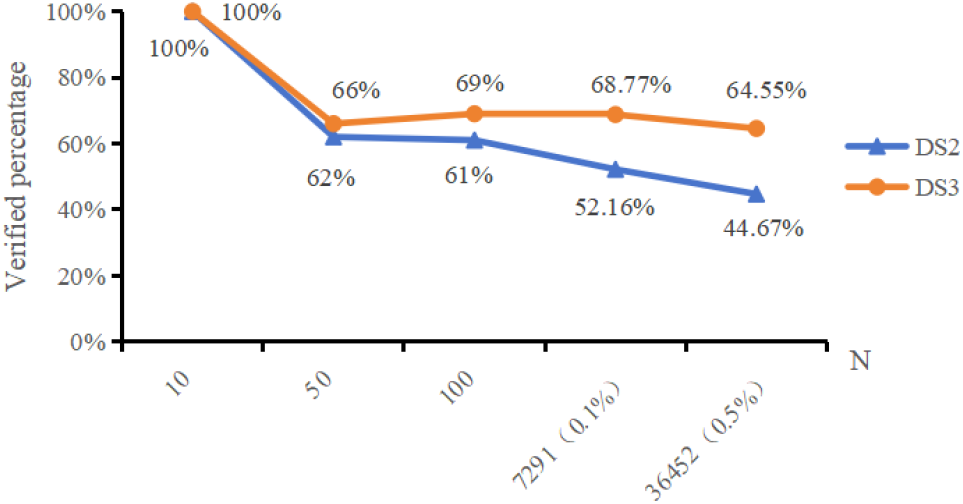
Result of version validation experiment

**Table 4.**
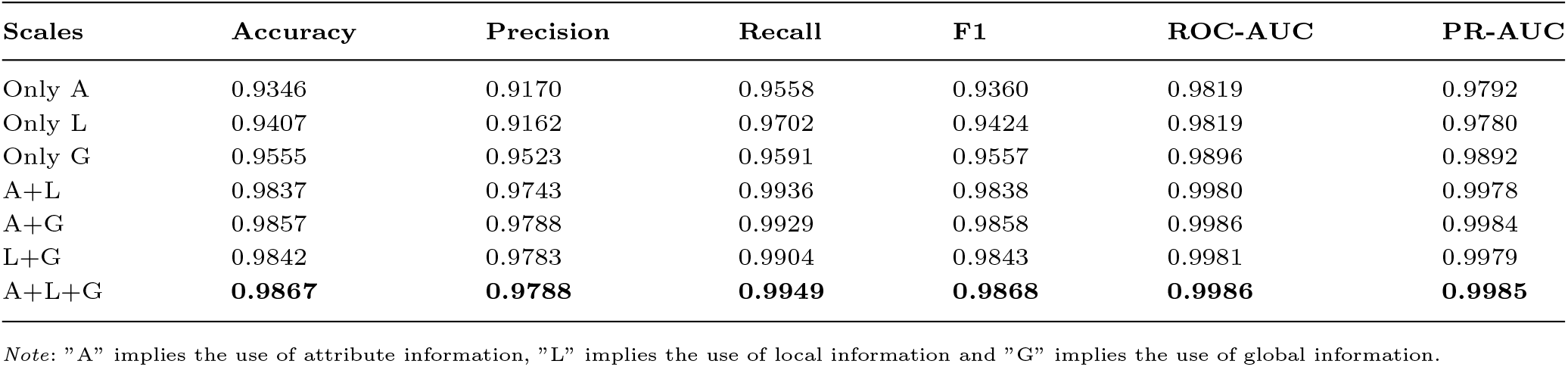
Results of ablation test on ALG–DDI with DS2.

**Table 5.** Results of comparing with state-of-the-art methods on Multi-DS.

| Model | Accuracy | macro-P | macro-R | macro-F1 | AUROC | AUPR |
| --- | --- | --- | --- | --- | --- | --- |
| RF | 0.7775 | 0.7893 | 0.5161 | 0.5936 | 0.9956 | 0.8349 |
| LR | 0.7920 | 0.7437 | 0.5236 | 0.5948 | 0.9960 | 0.8400 |
| DNN | 0.8797 | 0.8047 | 0.7027 | 0.7223 | 0.9963 | 0.9134 |
| DeepDDI | 0.8371 | 0.7275 | 0.6611 | 0.6848 | 0.9961 | 0.8899 |
| DDIMDL | 0.8852 | 0.8471 | 0.7182 | 0.7585 | 0.9976 | 0.9208 |
| Lee et al | 0.9094 | 0.8509 | 0.8339 | 0.8391 | 0.9961 | 0.9562 |
| **ALG-DDI** | **0.9142** | **0.8901** | **0.8449** | **0.8579** | **0.9964** | **0.9637** |

**Table 6.** Top 15 Predicted Drugs.

| Rank | Drug Name | DDI Description |
| --- | --- | --- |
| 1 | Desmopressin | The risk or severity of hypertension can be increased when Desmopressin is combined with Cannabidiol. |
| 2 | Cyclosporine | The serum concentration of Cyclosporine can be increased when it is combined with Cannabidiol. |
| 3 | Folic acid | Cannabidiol may decrease the excretion rate of Folic acid which could result in a higher serum level. |
| 4 | Fluvoxamine | The risk or severity of adverse effects can be increased when Cannabidiol is combined with Fluvoxamine. |
| 5 | Valsartan | The metabolism of Cannabidiol can be decreased when combined with Valsartan. |
| 6 | Amphetamine | The metabolism of Amphetamine can be decreased when combined with Cannabidiol. |
| 7 | Nicotine | The metabolism of Nicotine can be decreased when combined with Cannabidiol. |
| 8 | Cevimeline | The metabolism of Cevimeline can be decreased when combined with Cannabidiol. |
| 9 | Lorazepam | The risk or severity of adverse effects can be increased when Cannabidiol is combined with Lorazepam. |
| 10 | Goserelin | N.A. |
| 11 | Esmolol | The metabolism of Esmolol can be decreased when combined with Cannabidiol. |
| 12 | Bortezomib | The serum concentration of Bortezomib can be increased when it is combined with Cannabidiol. |
| 13 | Tramadol | The risk or severity of serotonin syndrome can be increased when Cannabidiol is combined with Tramadol. |
| 14 | Betaxolol | The metabolism of Betaxolol can be decreased when combined with Cannabidiol. |
| 15 | Sildenafil | The metabolism of Sildenafil can be decreased when combined with Cannabidiol. |

### Ablation experiments

To validate the necessity and contribution degree of information at each scale, we designed six different variants for the ablation study, with three of them using single-scale drug information for experimentation, and the other three using information from dual-scale information, The results are presented in 6.

Firstly, when we use information from all scales, the model achieves the best performance. Reducing any of the scales results in a performance drop, and employing dual-scale information is more effective than using a single scale alone. These findings strongly demonstrate the necessity and effectiveness of multi-scale feature fusion, indicating that ALG-DDI can indeed enhance the performance through multiple information complementing.

Additionally, we observed that in single-scale experiments, using global information (G) yielded the best results, and in dual-scale experiments, the performance drop was most significant when not using only global information compared to using all scales (A+L+G). This implies that the impact of global information extracted from the knowledge graph is greater compared to other scales. It indicates the correctness of introducing the knowledge graph into the DDI prediction task.

### Model performance on DDI events task

We tested the performance of ALG-DDI on DDI event prediction task, the evaluating metrics include *macro − P*, *macro − R, macro − F* 1, Accuracy, AUROC, and AUPR. We compared ALG-DDI with other state-of-the-art DDI event prediction models, including DeepDDI [4], DDIMDL [37], and Lee et al [38]. Additionally, we also compared ALG-DDI with general classification methods, such as Random Forest (RF), Deep Neural Network (DNN), and Logistic Regression (LR). The average results of 5-fold cross-validation under the same experimental conditions are presented in Table 5.

The results show that the top 10 predicted DDIs have been successively confirmed by subsequent researchers. Even among the top 0.5% (i.e., the top 36452), 44.67% of them were confirmed in DS2 (published in 2017), and 64.55% of them were confirmed in DS3 (published in 2022). This further illustrates that ALG-DDI possesses good foresight and have a excellent performance of potential DDI identification.

### Case study:Cannabidiol

To validate the excellent performance of ALG-DDI in practical prediction tasks, we conducted a case study using Cannabidiol(CBD). Specifically, we trained the model with the latest DS3 dataset and then scored all unknown drug pairs involving Cannabidiol that were not present in DS3. Higher scores indicate a greater likelihood of a relationship with Cannabidiol. The top 10 predicted drugs are shown in **??**, where nine of them have been further confirmed and included in the DrugBank database, along with detailed DDI descriptions.

To provide a biomedical speculation for the potential interaction between unconfirmed Cannabidiol-Goserelin, we consulted their pathophysiological knowledge. Cannabidiol have anti-tumour effects, which induce cell death pathways, cell growth arrest and tumour angiogenesis invasion and metastasis inhibition [39].Goserelin is a gonadotropin-releasing hormone (GnRH) analogue frequently utilized in the management of hormone-related tumors, including breast cancer and prostate cancer [40].Although their mechanisms of action differ, both can act on the relevant symptoms of breast cancer, and in certain situations, they may impact aspects like cancer cell growth, inflammatory responses, or cell apoptosis through different pathways.The relevant study by Alsherbiny et al. [41] also suggests that the cytotoxic mechanisms of CBD are encouraging, providing impetus for further research on the interaction between CBD and hormonal therapies, including aromatase inhibitors and new-generation drugs such as goserelin.

## Conclusions

In this article, we introduced a model named ALG-DDI, which leverages a Transformer encoder to integrate multi-scale drug information for DDI prediction tasks, including attribute information from molecular graphs, local information from heterogeneous graphs, and global information from knowledge graphs. Experimental results have proved that our proposed model is better than the state-of-the-art models. In addition, we demonstrated the effectiveness and robustness of our model by conducting experiments across various datasets and employing different fusion strategies. The ablation study also demonstrated the necessity of each scale in feature fusion. In summary, ALG-DDI is an effective model for discovering potential DDIs which can be used for preliminary screening of potential DDIs before the wet laboratory experiments. In the future work, we aim to further optimize the model and bring the idea of multi-scale drug information fusion to the DDI events prediction task, with the expectation of achieving good results.

## References

1. Supinya Dechanont, Sirada Maphanta, Bodin Butthum, and Chuenjid Kongkaew. Hospital admissions/visits associated with drug–drug interactions: a systematic review and meta-analysis. Pharmacoepidemiology and drug safety, 23(5):489–497, 2014.

2. Solène M Laville, Val’ serie Gras-Champel, Julien Moragny, Marie Metzger, Christian Jacquelinet, Christian Combe, Denis Fouque, Maurice Laville, Luc Frimat, Bruce M Robinson, et al. Adverse drug reactions in patients with ckd. Clinical journal of the American Society of Nephrology: CJASN, 15(8):1090, 2020.

3. Santiago Vilar, Rave Harpaz, Eugenio Uriarte, Lourdes Santana, Raul Rabadan, and Carol Friedman. Drug—drug interaction through molecular structure similarity analysis. Journal of the American Medical Informatics Association, 19(6):1066–1074, 2012.

4. Jae Yong Ryu, Hyun Uk Kim, and Sang Yup Lee. Deep learning improves prediction of drug–drug and drug–food interactions. Proceedings of the national academy of sciences, 115(18):E4304–E4311, 2018.

5. Wen Zhang, Yanlin Chen, Feng Liu, Fei Luo, Gang Tian, and Xiaohong Li. Predicting potential drug-drug interactions by integrating chemical, biological, phenotypic and network data. BMC bioinformatics, 18:1–12, 2017.

6. Sheng Qian, Siqi Liang, and Haiyuan Yu. Leveraging genetic interactions for adverse drug-drug interaction prediction. PLoS computational biology, 15(5):e1007068, 2019.

7. Assaf Gottlieb, Gideon Y Stein, Yoram Oron, Eytan Ruppin, and Roded Sharan. Indi: a computational framework for inferring drug interactions and their associated recommendations. Molecular systems biology, 8(1):592, 2012.

8. Feixiong Cheng and Zhongming Zhao. Machine learning-based prediction of drug–drug interactions by integrating drug phenotypic, therapeutic, chemical, and genomic properties. Journal of the American Medical Informatics Association, 21(e2):e278–e286, 2014.

9. Narjes Rohani and Changiz Eslahchi. Drug-drug interaction predicting by neural network using integrated similarity. Scientific reports, 9(1):13645, 2019.

10. Aurel Cami, Shannon Manzi, Alana Arnold, and Ben Y Reis. Pharmacointeraction network models predict unknown drug-drug interactions. PloS one, 8(4):e61468, 2013.

11. Marinka Zitnik, Monica Agrawal, and Jure Leskovec. Modeling polypharmacy side effects with graph convolutional networks. Bioinformatics, 34(13):i457–i466, 2018.

12. Md Rezaul Karim, Michael Cochez, Joao Bosco Jares, Mamtaz Uddin, Oya Beyan, and Stefan Decker. Drug-drug interaction prediction based on knowledge graph embeddings and convolutional-lstm network. In Proceedings of the 10th ACM international conference on bioinformatics, computational biology and health informatics, `pages 113–123, 2019.

13. Yue-Hua Feng, Shao-Wu Zhang, and Jian-Yu Shi. Dpddi: a deep predictor for drug-drug interactions. BMC Bioinformatics, 21(1):1–15,2020.

14. Fei Wang, Xiujuan Lei, Bo Liao, and Fang-Xiang Wu. Predicting drug–drug interactions by graph convolutional network with multi-kernel. Briefings in Bioinformatics, 23(1):bbab511, 2022.

15. Shenggeng Lin, Yanjing Wang, Lingfeng Zhang, Yanyi Chu, Yatong Liu, Yitian Fang, Mingming Jiang, Qiankun Wang, Bowen Zhao, Yi Xiong, et al. Mdf-sa-ddi: predicting drug– drug interaction events based on multi-source drug fusion, multi-source feature fusion and transformer self-attention mechanism. Briefings in Bioinformatics, 23(1):bbab421, 2022.

16. Hui Yu, Kui-Tao Mao, Jian-Yu Shi, Hua Huang, Zhi Chen, Kai Dong, and Siu-Ming Yiu. Predicting and understanding comprehensive drug-drug interactions via semi-nonnegative matrix factorization. BMC systems biology, 12(1):101–110, 2018.

17. Jiajing Zhu, Yongguo Liu, Yun Zhang, and Dongxiao Li. Attribute supervised probabilistic dependent matrix tri-factorization model for the prediction of adverse drug-drug interaction. IEEE Journal of Biomedical and Health Informatics, 25(7):2820–2832, 2020.

18. Vivian Law, Craig Knox, Yannick Djoumbou, Tim Jewison, An Chi Guo, Yifeng Liu, Adam Maciejewski, David Arndt, Michael Wilson, Vanessa Neveu, et al. Drugbank 4.0: shedding new light on drug metabolism. Nucleic acids research, 42(D1):D1091–D1097, 2014.

19. David S Wishart, Yannick D Feunang, An C Guo, Elvis J Lo, Ana Marcu, Jason R Grant, Tanvir Sajed, Daniel Johnson, Carin Li, Zinat Sayeeda, et al. Drugbank 5.0: a major update to the drugbank database for 2018. Nucleic acids research, 46(D1):D1074–D1082, 2018.

20. Payal Chandak, Kexin Huang, and Marinka Zitnik. Building a knowledge graph to enable precision medicine. Scientific Data, 10(1):67, 2023.

21. Greg Landrum. Rdkit documentation. Release, 1(1-79):4, 2013.

22. Weihua Hu, Bowen Liu, Joseph Gomes, Marinka Zitnik, Percy Liang, Vijay Pande, and Jure Leskovec. Strategies for pre-training graph neural networks. arXiv preprint arXiv:1905.12265, 2019.

23. Jie Zhou, Ganqu Cui, Shengding Hu, Zhengyan Zhang, Cheng Yang, Zhiyuan Liu, Lifeng Wang, Changcheng Li, and Maosong Sun. Graph neural networks: A review of methods and applications. AI open, 1:57–81, 2020.

24. Zhong-Hao Ren, Zhu-Hong You, Chang-Qing Yu, Li-Ping Li, Yong-Jian Guan, Lu-Xiang Guo, and Jie Pan. A biomedical knowledge graph-based method for drug– drug interactions prediction through combining local and global features with deep neural networks. Briefings in Bioinformatics, 23(5):bbac363, 2022.

25. Will Hamilton, Zhitao Ying, and Jure Leskovec. Inductive representation learning on large graphs. Advances in neural information processing systems, 30, 2017.

26. Michael Schlichtkrull, Thomas N Kipf, Peter Bloem, Rianne Van Den Berg, Ivan Titov, and Max Welling. Modeling relational data with graph convolutional networks. In The Semantic Web: 15th International Conference, ESWC 2018, Heraklion, Crete, Greece, June 3–7, 2018,

27. Matthias Fey and Jan Eric Lenssen. Fast graph representation learning with pytorch geometric. arXiv preprint arXiv:1903.02428, 2019.

28. Th’ seo Trouillon, Johannes Welbl, Sebastian Riedel, Eric Gaussier, and Guillaume Bouchard. Complex embeddings for simple link prediction. In International conference on machine learning, pages 2071–2080. PMLR, 2016.

29. Kaiming He, Xiangyu Zhang, Shaoqing Ren, and Jian Sun. Deep residual learning for image recognition. In Proceedings of the IEEE conference on computer vision and pattern recognition, pages 770–778, 2016.

30. Jimmy Lei Ba, Jamie Ryan Kiros, and Geoffrey E Hinton. Layer normalization. arXiv preprint arXiv:1607.06450, 2016.

31. Tsung-Yi Lin, Priya Goyal, Ross Girshick, Kaiming He, and Piotr Doll’ sar. Focal loss for dense object detection. In Proceedings of the IEEE international conference on computer vision, pages 2980–2988, 2017.

32. Xin Chen, Xien Liu, and Ji Wu. Gcn-bmp: investigating graph representation learning for ddi prediction task. Methods, 179:47–54, 2020.

33. Mengying Sun, Fei Wang, Olivier Elemento, and Jiayu Zhou. Structure-based drug-drug interaction detection via expressive graph convolutional networks and deep sets (student abstract). In proceedings of the AAAI conference on artificial intelligence, volume 34, pages 13927–13928, 2020.

34. Nuo Xu, Pinghui Wang, Long Chen, Jing Tao, and Junzhou Zhao. Mr-gnn: Multi-resolution and dual graph neural network for predicting structured entity interactions. arXiv preprint arXiv:1905.09558, 2019.

35. Xusheng Cao, Rui Fan, and Wanwen Zeng. Deepdrug: a general graph-based deep learning framework for drug relation prediction. biorxiv, 2020.

36. Arnold K Nyamabo, Hui Yu, and Jian-Yu Shi. Ssi–ddi: substructure–substructure interactions for drug–drug interaction prediction. Briefings in Bioinformatics, 22(6):bbab133, 2021.

37. Yifan Deng, Xinran Xu, Yang Qiu, Jingbo Xia, Wen Zhang, and Shichao Liu. A multimodal deep learning framework for predicting drug–drug interaction events. Bioinformatics, 36(15):4316–4322, 2020.

38. Geonhee Lee, Chihyun Park, and Jaegyoon Ahn. Novel deep learning model for more accurate prediction of drug-drug interaction effects. BMC bioinformatics, 20:1–8, 2019.

39. Dimakatso R. Mokoena, Blassan P. George, and Heidi Abrahamse. Enhancing breast cancer treatment using a combination of cannabidiol and gold nanoparticles for photodynamic therapy. International journal of molecular sciences, 20(19):4771, 2019.

40. Ian D Cockshott. Clinical pharmacokinetics of goserelin. Clinical pharmacokinetics, 39:27–48, 2000.

41. Muhammad A Alsherbiny, Deep J Bhuyan, Mitchell N Low, Dennis Chang, and Chun Guang Li. Synergistic interactions of cannabidiol with chemotherapeutic drugs in mcf7 cells: Mode of interaction and proteomics analysis of mechanisms. International journal of molecular sciences, 22(18):10103, 2021.

